# Postembryonic development of the two-spotted field cricket (*Gryllus bimaculatus*): a staging system

**DOI:** 10.1101/2021.02.24.432775

**Authors:** Jakke Neiro

## Abstract

The two-spotted field cricket *Gryllus bimaculatus* has emerged as a central model for studies on insect development, regeneration, and physiology. *G. bimaculatus* has the most sophisticated functional genetic toolkit of any hemimetabolous insect, making it a foremost model to understand the evolutionary developmental biology and comparative physiology of insects. However, the morphology and stages of postembryonic development have never been comprehensively reported. Here, 8 morphologically defined stages are described. Size, coloration, and the morphology of wing buds, hind tibial spines, and the ovipositor are the best landmarks for staging. The stages correspond to the 8-12 moult-based instars present in the literature. The staging system aims to standardise studies on the postembryonic development of *G. bimaculatus* and serve as a point of reference for delineating interspecific postembryonic homologies within Orthoptera.

## 1. Introduction

The two-spotted field cricket *Gryllus bimaculatus* has emerged as a central model for studies on insect development, regeneration, and physiology (Mito & Noji, 2008; Donoughe & Extavour, 2016; Horch et al., 2017; Ylla et al., 2020). *G. bimaculatus* has the most sophisticated functional genetic toolkit of any hemimetabolous insect, making it a foremost model to understand the evolutionary developmental biology (evo-devo) and comparative physiology of insects (Donoughe & Extavour, 2016). Furthermore, *G. bimaculatus* is consumed and farmed in different parts of the world and has the promise to become an important species in the global food industry (Hansboonsong et al., 2013; Sorjonen et al., 2019; Ylla et al, 2020).

The morphology of adult crickets represents well the general morphology of insects (Fig. 1 A to D). Surprisingly, although embryonic development has been well staged in *G. bimaculatus*, no morphological staging of the postembryonic development seems to have been completed (Donoughe & Extavour, 2016; Horch et al., 2017). Furthermore, different authors have reported variable values for the number of instars as a part of other studies on embryonic development or leg regeneration, ranging from 8 to 12 (Mito et al., 2002; Nakamura et al., 2007; Nakamura et al., 2008; Bando et al., 2009; Bando et al., 2011a; Bando et al., 2011b; Hamada et al., 2015; Ishimaru et al., 2015; Donoughe & Extavour 2016; Ishimaru et al., 2016; Mashimo & Machida, 2017; Ishimaru et al., 2018). Nonetheless, three primary experiments have been reported: Sturm (2002) counted the moults and found 8 to 10 instars depending on temperature, Suzuki and Nishimura (1997) determined 8 instars based on head width and the position of wing buds, while Zhemchuzhnikov and Knyazev (2012) cited Knyazev (1985) who defined 6 instars based on morphological landmarks. Most thoroughly, Jobin (1961) described the morphology of all the instars in the related *G. assimilis*, *G. rubens*, *G. veletis*, and *G. pennysylvanicus*, concluding that the number of instars varies from 8 to 12 depending on temperature and other environmental factors. This finding is in accordance with even earlier reports of *G. veletis*, *G. pennysylvanicus*, and *G. rubens*, which claimed that the instar number was somewhere between 8 and 12 (Criddle, 1925; Folsom, 1934; Severin, 1926; Severin, 1935).

**Fig. 1.**
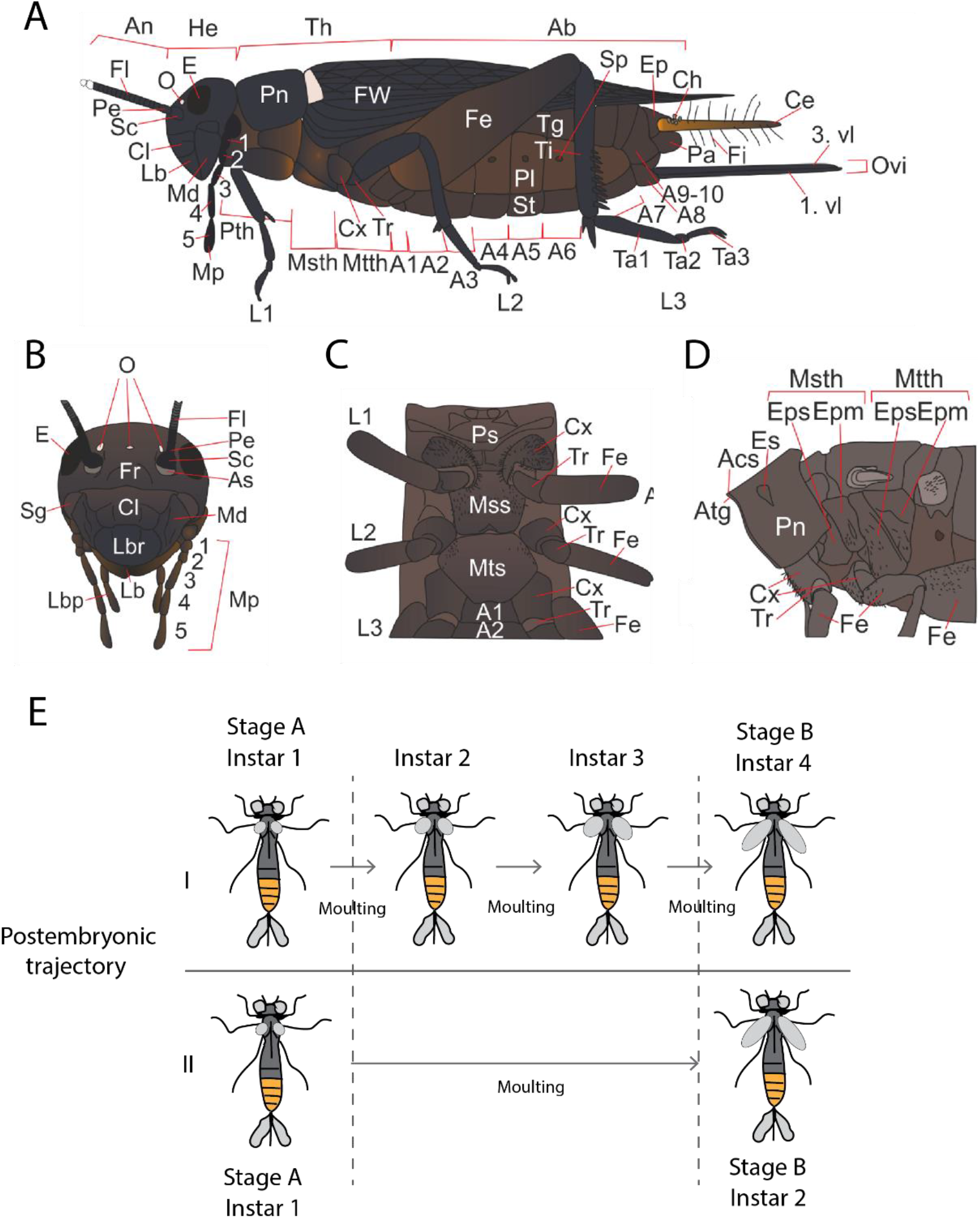
External morphology of the adult cricket and staging postembryonic development of insects. (A) Lateral view. (B) Anterior view of the head. (C) Ventral view of the thorax. (D) Lateral view of the thorax (wings removed). (E) Two postembryonic trajectories I and II are compared to contrast instars and stages. The stage (A and B) denotes a certain morphological state (small or big wing buds), while instar (1,2,3, and 4) denotes a period between two moults. As the moults may vary depending on environmental factors, the stages offer a standardised way to characterise postembryonic development in insects. Ab: Abdomen, Acs: Antecostal suture, An: Antenna, As: Antennal socket, Atg: Acrotergite, Ce: Cercus, Ch: Clavate hair, Cl: Clypeus, Cx: Coxa, E: Eye, Ep: Epiproct, Epm: Epimeron, Eps: Episternum, Es: Eyespot, Fe: Femur, Fi: Filiform hair, Fl: Flagellum, Fr: Frons, FW: Forewing, He: Head, L1-3: Leg 1-3, Lb: Labium, Lbp: Labial palp, Lbr: Labrum, Md: Mandible, Mp: Maxillary palp, Mss: Mesosternum, Msth: Mesothorax, Mts: Metasternum, Mtth: Metathorax, O: Ocellus, Ovi: Ovipositor, Pa: Paraproct, Pl: Pleuron, Pn: Pronotum, Ps: Prosternum, Pth: Prothorax, Scp: Scape, Sg: Subgena, Sp: Spiracle, St. Sternum, Ta1-3: Tarsus 1-3, Tg: Tergum, Ti: Tibia, Tr: Trochanter, 1. vl: 1st valvifer, 3. vl: 3rd valvifer.

Taken together, the morphology of the wing pads, the spines of the hind tibiae and the sexual characteristics of the abdomen are the most useful features to characterise instars from each other in the genus *Gryllus* (Criddle, 1925; Jobin, 1961; Severin, 1935). Multiple environmental factors are known to affect the number of instars in insects, and therefore the usual practice of staging postembryonic development based on moulting and instars is not optimal (Fig. 1 E) (Esperk et al., 2007; Minelli and Fusco, 2013). As advocated by Minelli and Fusco (2013), morphologically defined stages are to be preferred over moulting-based instars for periodisation of postembryonic development in arthropods (Fig. 1 E). Here, 8 morphologically defined stages are described for the postembryonic development of *G. bimaculatus*.

## 2. Materials and methods

### 2.1 Animal husbandry

The laboratory stocks of *G. bimaculatus* were purchased from ReptileManiacs (ReptileManiacs Ay, Y-tunnus: 2440893-2, Tampere, Finland). The nymphal stages were maintained in medium-sized plastic boxes with a perforated lid (28 cm × 18 cm × 12.5 cm), while the adults were held in bigger terraria (37 cm × 22 cm × 18 cm). Crampled paper was provided to reduce the risk of cannibalism. The crickets were fed JBL NovoBel Flakes, and water was provided every second day by moistening tissue paper. The eggs were collected from the adult terrarium by collecting the tissue paper and incubating it in a small plastic box (13.5 cm × 10 cm × 7 cm) until hatching. The tissue paper bearing the eggs was moistened each day to keep the eggs hydrated. The crickets and the eggs were maintained at 30 °C under 12L-12D light conditions.

### 2.2 Staging postembryonic development

The life cycle of *G. bimaculatus* was followed daily for 28 days. The nymphal stages were imaged with a stereomicroscope (Leica EZ4W) using the Leica LAS EZ (v 3.4) software. Maximal illumination (the first illumination option) was applied. The imaged nymphs were freezed in − 20 °C, after which they were mounted on a dissection pin attached to a piece of styrox foam. White balance was adjusted by having a sheet of white paper as the reference. When imaging hind tibiae, the grey stage plate of the microscope was set as the background. Images were taken manually from different focal planes, which were stacked together using the CombineZP (v 1.0) software. Focal stacking was performed by selecting "All methods". The images were cropped and annotated in CorelDRAW 2017.

## 3. Results

### 3.1 Stage 1

At the first stage, the total body length from prostomium to periproct is about 2.5 − 3.5 mm. The head (cephalon) is hemispherical in shape, always shiny black and wider than the prothorax (Fig. 2 A and C). The lateral ocelli are rather indistinct but nevertheless visible as white to light brown dots (Fig. 2 A and D). The anterior pronotal margin has a row of ten black setae, five on each side of the midline, protruding forward to form a bridge-like fringe over the neck, while the posterior margin has 10 setae protruding straight backward (Fig. 2 A and D). The medial dorsal ecdysial suture is visible as a narrow light-yellow line in the middle of the pronotum (Fig. 2 D). The posterior mesonotal margin has also a fringe of ten black setae (five on each side), protruding backward (Fig. 2 A). The metanotum is shiny black and has a fringe of a variable number of black setae directed backward at the posterior margin (Fig. 2 A). The pterothoracic sclerites (episternum and epimeron) are brownish black, small, round, and only lightly sclerotised (Fig. 2 A and D). The pleural membrane is greyish brown (Fig. 2 B). The thoracic sternites are dark brown to shiny black (Fig. 2 C). The basal part of cerci is yellow to orange, and the tip is grey to black (Fig. 2 B and D). Cerci bear long, protrusive reddish grey filiform hairs and small black hairs. At the base of cerci, one or two clavate hairs (bulbous sensilla) are occasionally present (Fig. 2 B and D). The femora and tibiae of all legs are shiny black, and the hindleg bears no tibial spines (Fig. 2 D). No external sexual characters are apparent (Fig. 2 B).

**Fig. 2.**
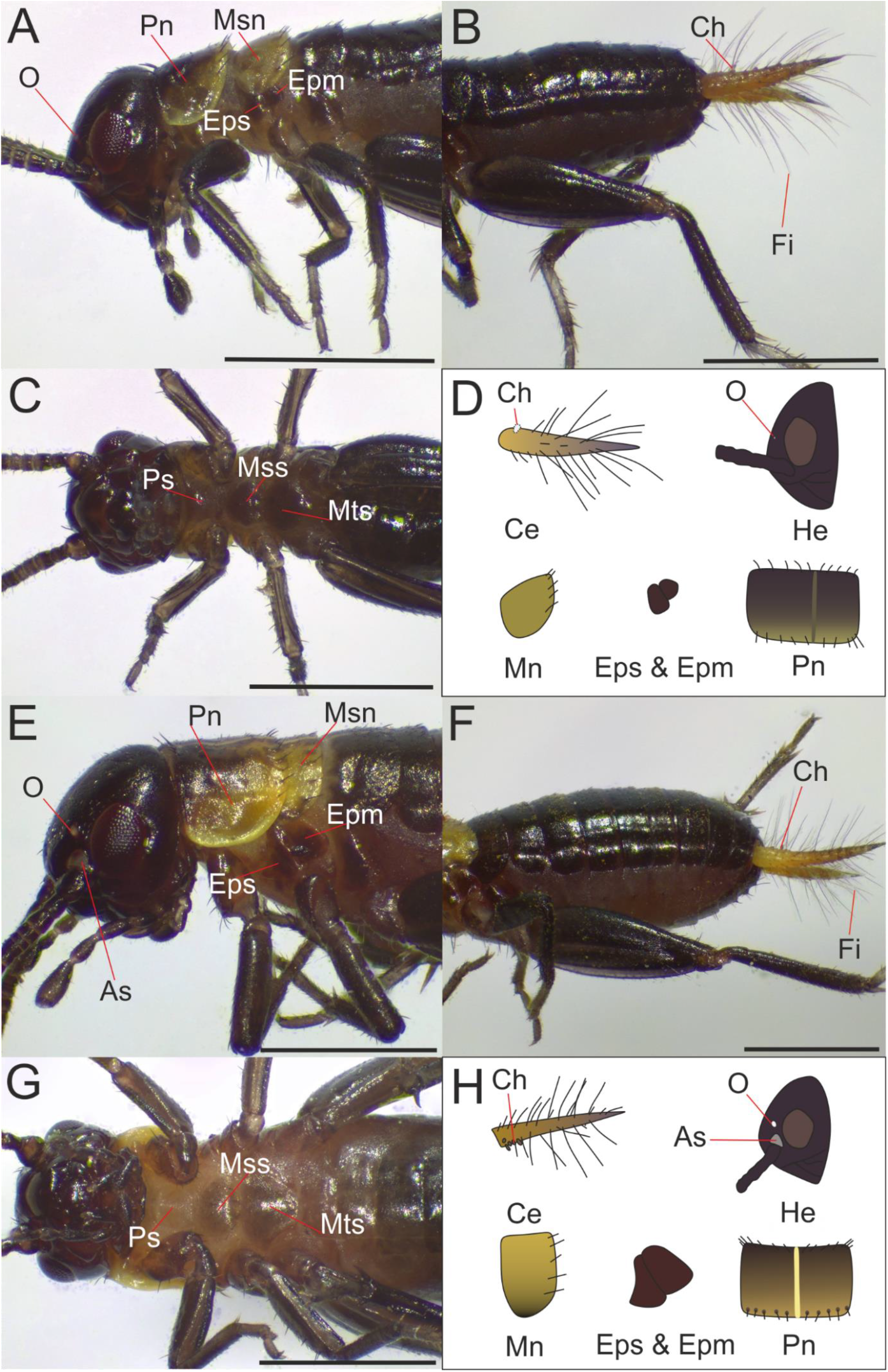
External morphology of the first and second stages. (A) Lateral view of the head and thorax of the first stage. (B) Lateral view of the abdomen of the first stage. (C) Ventral view of the head and thorax of the first stage. (D) Morphological characteristics of the first stage: none to one clavate hair, indistinct lateral ocelli, uniformly yellowish brown mesonotum, small and round episternum and epimeron, and 10-12 curly black setae at the anterior end of the pronotum. (E) Lateral view of the head and thorax of the second stage. (F) Lateral view of the abdomen of the second stage. (G) Ventral view of the head and thorax of the second stage (H) Morphological characteristics of the second stage: two to six clavate hairs, distinct lateral ocelli and antennal socket, pigmented margin of the mesonotum, robust and wedge-formed episternum and epimeron, 14-16 straight black setae at the anterior end of the pronotum, and a bright ecdysial suture dividing the pronotum. As: Antennal socket, Ce: Cercus, Ch: Clavate hair, Epm: Epimeron, Eps: Episternum, Fi: Filiform hair, Msn: Mesonotum, Mss: Mesosternum, Mts: Metasternum, O: Ocellus, Ovi: Ovipositor, Pn: Pronotum, Ps: Prosternum.

### 3.2 Stage 2

At the second stage, the total body length from prostomium to periproct is 3.5 − 5.0 mm. The head is somewhat rounder in appearance than at the first stage, and it is only slightly wider than the prothorax (Fig. 2 E and G). The ocelli are clearly white, and the antennal socket is clearly visible as a grey membrane against the shiny black head capsule (Fig. 2 E and H). The pronotum is slightly grooved at the anterior margin (Fig. 2 E and H). The lateral pronotal margin is brightly yellow and forms a conspicuous ridge (Fig. 2 E). The anterior margin has a row of 14 or 16 black setae, protruding straight forward and not forming a bridge-like fringe as at the first stage (Fig. 2 E and H). The medial dorsal ecdysial suture is clearly visible as a narrow yellow line running through the middle of the dorsal part of the thorax (Fig. 2 H). The thoracic pleurites and the pleural membrane are clearly brown as compared with the first stage (Fig. 2 E and F). The mesonotal epimeron is highly sclerotised and forms a wedge-like structure pointing posteriorly (Fig. 2 E and H). At the base of cerci, three to five clavate hairs are present in contrast to the first stage (Fig. 2 F and H). The pleural membrane is clearly brown (Fig. E and F). No tibial spines and external sexual characters are apparent (Fig. 2 F).

### 3.3 Stage 3

At the third stage, the total body length from prostomium to periproct is 5.0 − 8.0 mm. The head is as wide as the prothorax, the ocelli are clearly white, and the antennal socket is light grey (Fig. 3 A and C). The pronotum is as wide as the head and grooved at the anterior margin but covered with two eyespot-like markings and browner in color than in previous stages (Fig. 3 A and D). The anterior margin has a row of 20 to 22 black setae, and the ecdysial suture is clearly visible in the dorsal part of the thorax (Fig. 3 A and D). The wing bud is lateral and always brownish black in color (Fig. 3 A). The metanotal epimeron is slightly more sclerotised than in the second stage (Fig. 3 A and D). The prosternum is light brown to light grey, and the ecdysial sutures surrounding the prosternum form a rhombus-like figure (Fig. 3 C). The trochantin and precoxale are sclerotised and can be seen as lateral ridges on the ventral side of the mesothorax in contrast to the previous stage (Fig. 3 A and C). Abdominal terga are dark brown with a variable amount of black pigmentation (Fig. 3 B). The hind tibiae bear one to two pairs of spines, and occasionally a third outer spine is present (Fig. 3 B and D). Generally, no external sexual characters are present, but occasionally two small pairs of ovipositor primordia are protruding between the sternites (Fig. 3 D).

**Fig. 3.**
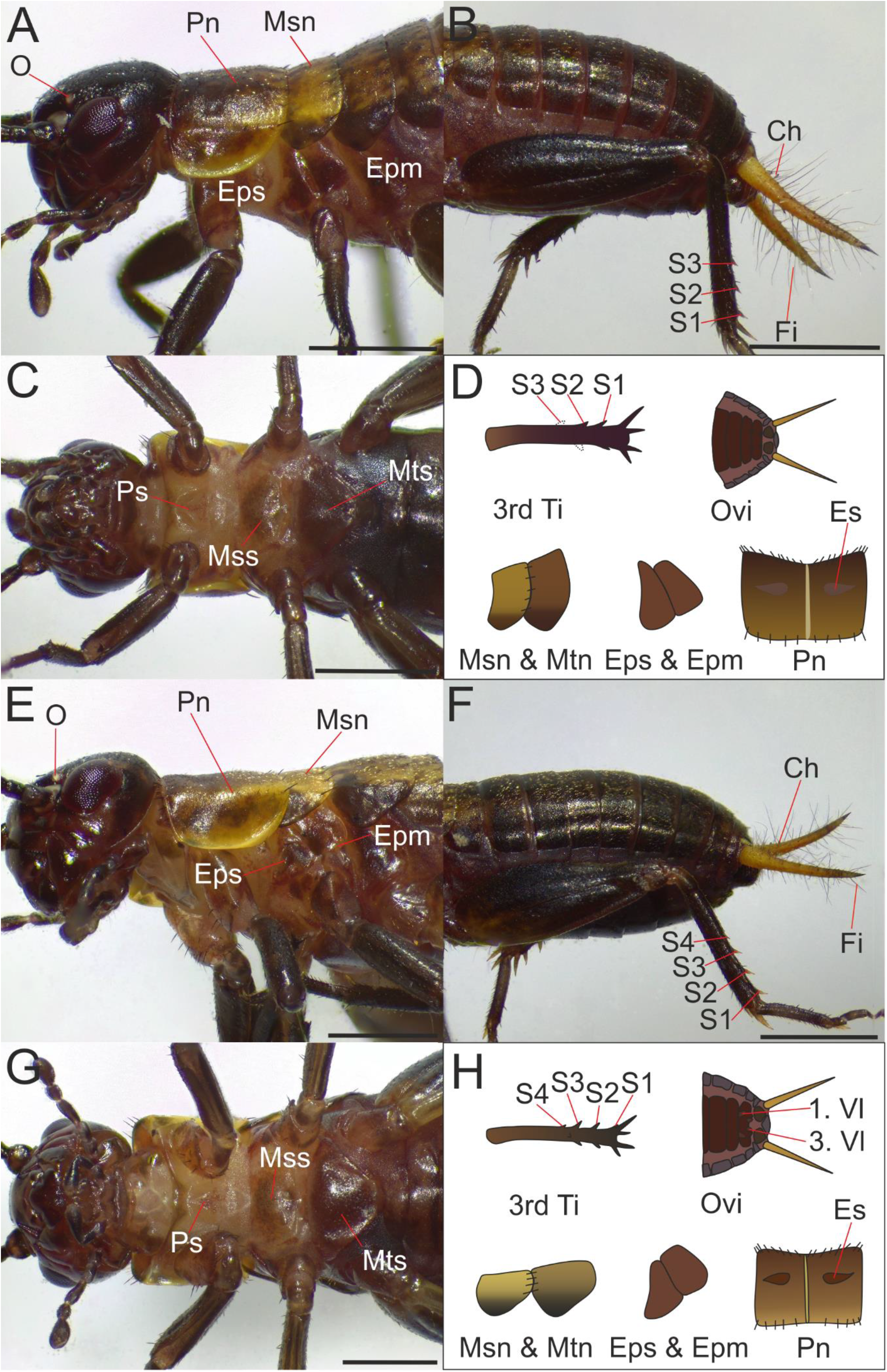
External morphology of the third and fourth stages. (A) Lateral view of the head and thorax of the third stage. (B) Lateral view of the abdomen of the third stage. (C) Ventral view of the head and thorax of the third stage. (D) Morphological characteristics of the third stage: two to three pairs of hind tibial spines, none or small rudimentary protrusions of valvifers, distinct lateral wing buds, light ecdysial suture in the pronotum and vague eyespot-like markings. (E) Lateral view of the head and thorax of the fourth stage. (F) Lateral view of the abdomen of the fourth stage. (G) Ventral view of the head and thorax of the fourth stage (H) Morphological characteristics of the fourth stage: three to four pairs of hind tibial spines, distinct valvifers, wedge-like wing buds, and distinct eyespot-like markings in the pronotum. Ch: Clavate hair, Epm: Epimeron, Eps: Episternum, Es: Eyespot-like marking, Fi: Filiform hair, Msn: Mesonotum, Mss: Mesosternum, Mtn: Metanotum, Mts: Metasternum, O: Ocellus, Ovi: Ovipositor, Pn: Pronotum, Ps: Prosternum, S: Spine, Vl: Valvifer.

### 3.4 Stage 4

At the fourth stage, the total body length from prostomium to periproct is 5.0 − 8.0 mm. The pronotum is as wide as the head as in the previous stage, but the anterior margin is slightly less grooved (Fig. 3 E, G and H). The coloration varies from black to dark and light brown (Fig. 3 E and H). Two eyespot-like cuticular markings are evident on each side of the dorsal midline (Fig. 3 H). The lateral pronotal margin is clearly yellow, although not as bright as in previous stages (Fig. 3 E). The wing buds are dark but more robust and wedge-like than in the previous stage (Fig. 3 E and H). The posterior tip of the mesonotal epimeron is more rounded than in the third stage (Fig. 3 E and H). A rhombus-like marking is visible in the ventral part of the neck (Fig. 3 G). The metasternum has a more robust posterior ridge than in previous stages (Fig. 3 G). Three pairs of tibial spines are evident and sometimes a fourth outer spine is visible (Fig. 3 F and H). The valvulae of the ovipositor are visible as two pairs of small buds in the eighth and ninth segments (Fig. 3 H).

### 3.5 Stage 5

At the fifth stage, the total body length from prostomium to periproct is 8.0 − 11.0 mm. The head is as wide as the prothorax, and the coloration varies from shiny black to dark brown (Fig. 4 A and C). The color of the pronotum varies from black to dark and light brown and is generally hairier than in previous stages (Fig. 4 A). The contour of the eyespot-like markings is even sharper than in the previous stage (Fig. 4 A). In contrast to previous stages, the anterior margin of the pronotum is covered in small brown to light brown hairs in addition to the dark setae (Fig. 4 D). The coloration of the mesonotum varies from dark brown to brownish yellow with black and brown patches, while the wing bud is always dark in color as in previous stages, but with a somewhat more rounded tip (Fig. 4 A and D). The mesonotal epimeron is more rectangular in shape than in the previous stage (Fig. 4 A and D). The prosternum is light brown to light grey, but the rhombus-like marking is less evident than in the fourth stage (Fig. 4 C). In contrast, the rhombus-like marking in the ventral part of the neck is as evident as in the fourth stage (Fig. 4 C). The prothoracic coxae are more robust than in the previous stage (Fig. 4 C). The mesosternum is dark brown and hairier than in previous stages (Fig. 4 C). The metasternum is wider and even more robust than in the fourth stage (Fig. 4 C). Abdominal terga are dark brown with a variable amount of black pigmentation (Fig. 4 B). There are four pairs of tibial spines, and occasionally a fifth outer spine is present (Fig. 4 B and D). The valvulae of the ovipositor are visible as two pairs of elongate buds in the eighth and ninth segments (Fig. 4 D).

**Fig. 4.**
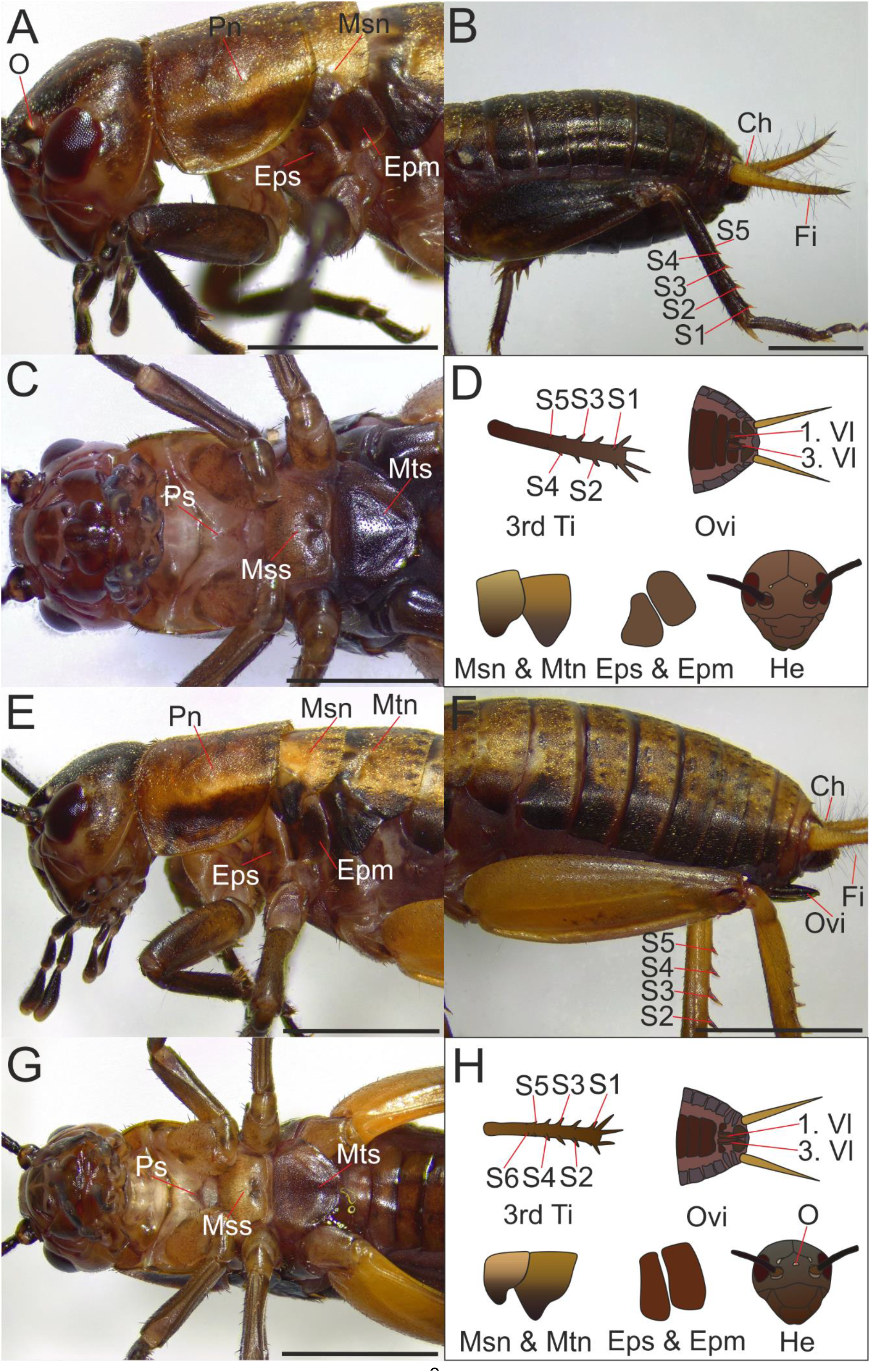
External morphology of the fifth and sixth stages. (A) Lateral view of the head and thorax of the fifth stage. (B) Lateral view of the abdomen of the fifth stage. (C) Ventral view of the head and thorax of the fifth stage. (D) Morphological characteristics of the fifth stage: five pairs of hind tibial spines, elongate valvifers, wing buds with a slightly grooved base, and no medial ocellus. (E) Lateral view of the head and thorax of the sixth stage. (F) Lateral view of the abdomen of the sixth stage. (G) Ventral view of the head and thorax of the sixth stage (H) Morphological characteristics of the sixth stage: six pairs of hind tibial spines, apposed valvifers, deep groove at the wing bud base, and medial ocellus. Ch: Clavate hair, Epm: Epimeron, Eps: Episternum, Es: Eyespot-like marking, Fi: Filiform hair, Msn: Mesonotum, Mss: Mesosternum, Mtn: Metanotum, Mts: Metasternum, O: Ocellus, Ovi: Ovipositor, Pn: Pronotum, Ps: Prosternum, S: Spine, Vl: Valvifer.

### 3.6 Stage 6

At the sixth stage, the total body length from prostomium to periproct is 12.0 − 15.0 mm. The medial ocellus is visible for the first time (Fig. 4 H). The pronotum is as wide as the head as in the third, fourth and fifth stages, but the anterior end is clearly narrower than the posterior end (Fig. 4 E). The color varies widely from black to dark and light brown (Fig. 4 E). The lateral pronotal margin is darker than in previous stages (Fig. 4 E). The medial dorsal ecdysial suture is clearly visible, but in contrast to previous stages, the line extends to the abdominal terga. The mesonotal wing bud is dark but has a deeper groove at the base than in the fifth stage (Fig. 4 E and H). The mesonotal epimeron is slightly narrower in shape than in the fifth stage (Fig. 4 E and H). The prosternum is light brown to light grey, but no rhombus-like marking visible as in the fourth and fifth stages (Fig. 4 G). Instead, a wedge-like sclerotised marking is apparent (Fig. 4 G). Similarly, the rhombus-like marking is missing from the ventral part of the neck (Fig. 4 G). There is almost no sternal membrane left, as the whole sternum is sclerotised (Fig. 4 G). In the femora and tibiae, coloration varies from dark brown to yellowish brown, and five to six pairs of tibial spines are present (Fig. 4 F and H). The valvulae appose to each other and form a distinct ovipositor that extends to the periproct, and the subgenital plate has developed into a characteristic triangular structure (Fig. 4 F and H).

### 3.7 Stage 7

At the seventh stage, the total body length from prostomium to periproct is 15.0 − 18.0 mm. The medial ocellus is even more apparent in the middle of the frontal cuticle than in the previous stage. The pro-, meso- and metanota vary widely from black to dark and light brown (Fig. 5 A and B). There are distinct sclerotised dots and margins in the ventral side of the neck (Fig. 5 A). The wing buds have turned into a dorsal position but are still black or dark brown in color (Fig. 5 A, B and D). The thoracic pleurites are dark to light brown, but almost no pleural membrane is visible as the pleuron is almost fully sclerotised (Fig. 5 A). The episterna and epimera have more or less the same shape as in the previous stage (Fig. 5 A). There are five to six tibial spines present as in the sixth stage (Fig. 5 D). The ovipositor extends slightly longer than the periproct (Fig. 5 B and D).

**Fig 5.**
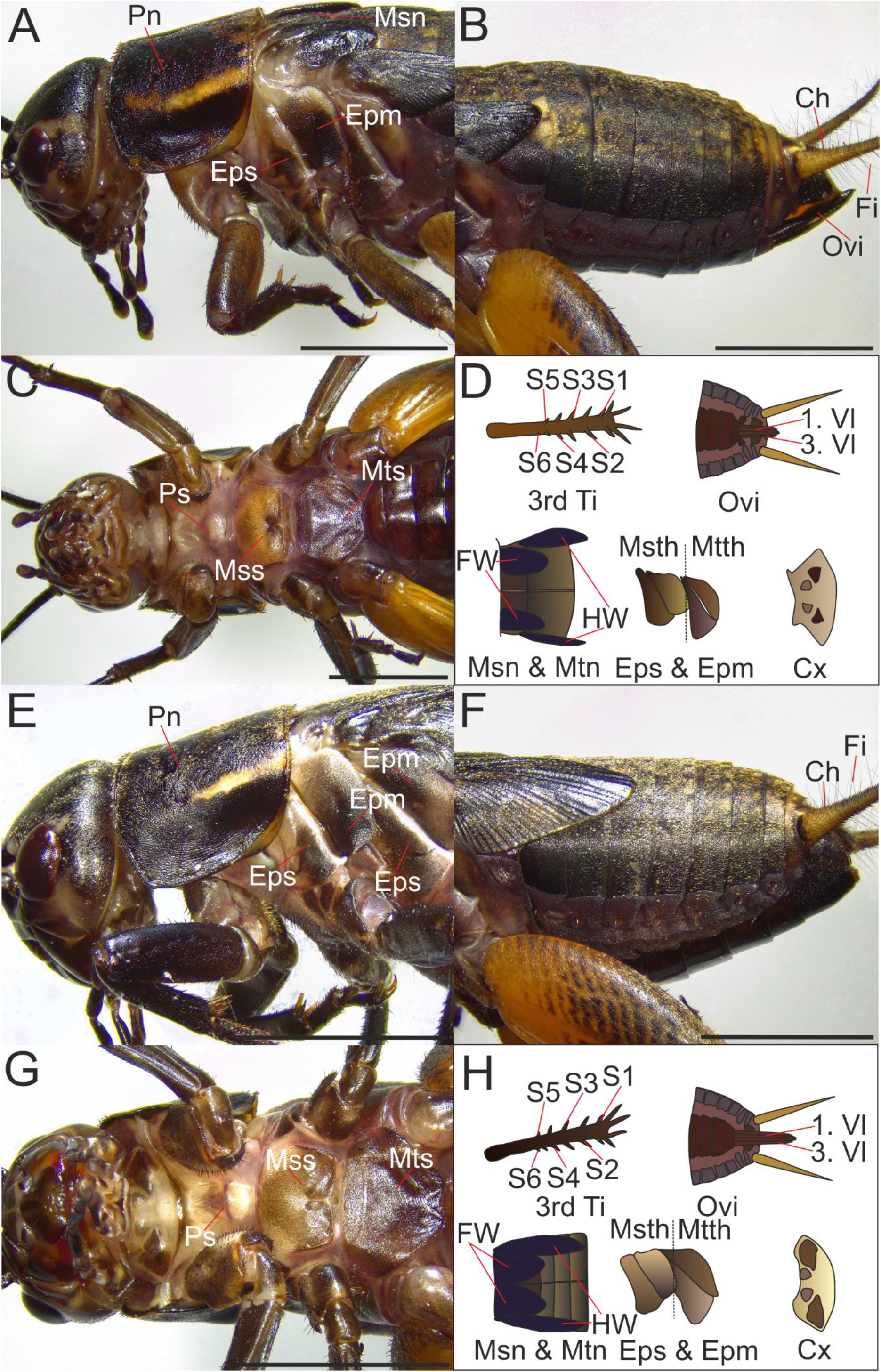
External morphology of the seventh and eighth stages. (A) Lateral view of the head and thorax of the seventh stage. (B) Lateral view of the abdomen of the seventh stage. (C) Ventral view of the head and thorax of the seventh stage. (D) Morphological characteristics of the seventh stage: six pairs of hind tibial spines with the sixth pair of spines small, ovipositor slightly longer than the periproct, hindwings extend just over the second segment, and small lateral sclerotised markings on the ventral side of the neck. (E) Lateral view of the head and thorax of the eighth stage. (F) Lateral view of the abdomen of the eighth stage. (G) Ventral view of the head and thorax of the eighth stage (H) Morphological characteristics of the eighth stage: six pairs of hind tibial spines with the sixth spine clerarly protrusive, ovipositor extends well over the periproct, hindwings extend to the fifth segment, and the rather big lateral sclerotised markings on the ventral side of the neck. Ch: Clavate hair, Cx: Cervix, Epm: Epimeron, Eps: Episternum, Es: Eyespot-like marking, Fi: Filiform hair, FW: Forewing, HW: Hindwing, Msn: Mesonotum, Mss: Mesosternum, Msth: Mesothorax, Mtn: Metanotum, Mts: Metasternum, Mtth: Metathorax, O: Ocellus, Ovi: Ovipositor, Pn: Pronotum, Ps: Prosternum, S: Spine, Vl: Valvifer.

### 3.8 Stage 8

At the eight stage, the total body length from prostomium to periproct is 18.0 − 23.0 mm. The wing buds are distinctly larger than in the seventh stage (Fig. 5 E, F and H). The lateral margins of the ventral part of the neck are more sclerotised and larger than in the previous stage (Fig. 5 G and H). The metathoracic epimeron is sclerotised and significantly larger than in previous stages (Fig. 5 E and H). About ten to fifteen clavate hairs are present at the base of the cerci (Fig. 5 F). Five to six pairs of tibial spines are present as in the previous stage (Fig. 5 H). The ovipositor extends longer than the periproct (Fig. 5 H).

## 4. Discussion

### 4.1 Staging of postembryonic development

The number and morphology of instars observed in *G. bimaculatus* is consistent with earlier works on *G. bimaculatus*, *G. assimilis*, *G. rubens*, *G. veletis*, and *G. pennsylvanicus* (Jobin, 1961; Mashimo & Machida, 2017). Thus, 8 morphologically identifiable stages can be defined, while the number of instars vary from 8 to 12 so that the sixth stage encompasses one instar at 30 °C and up to five instars at lower temperatures. In other words, the nymphs of *G. bimaculatus* do not advance morphologically between the 6th and 10th instars despite moulting. The nymphal stages can be identified based on size, coloration, and mainly the morphology of the wings, the hind tibia, and the ovipositor, as reported by Jobin (1961) (Table 1).

**Table 1.**
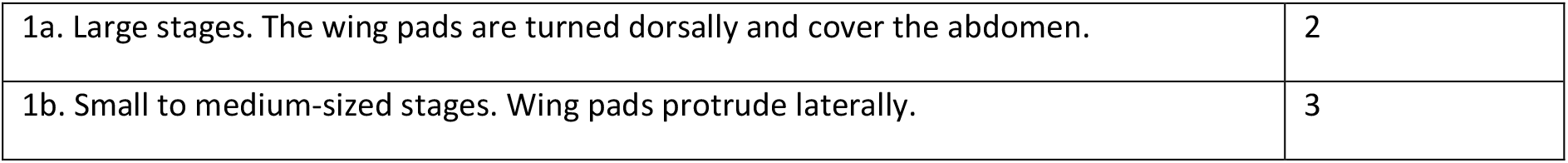
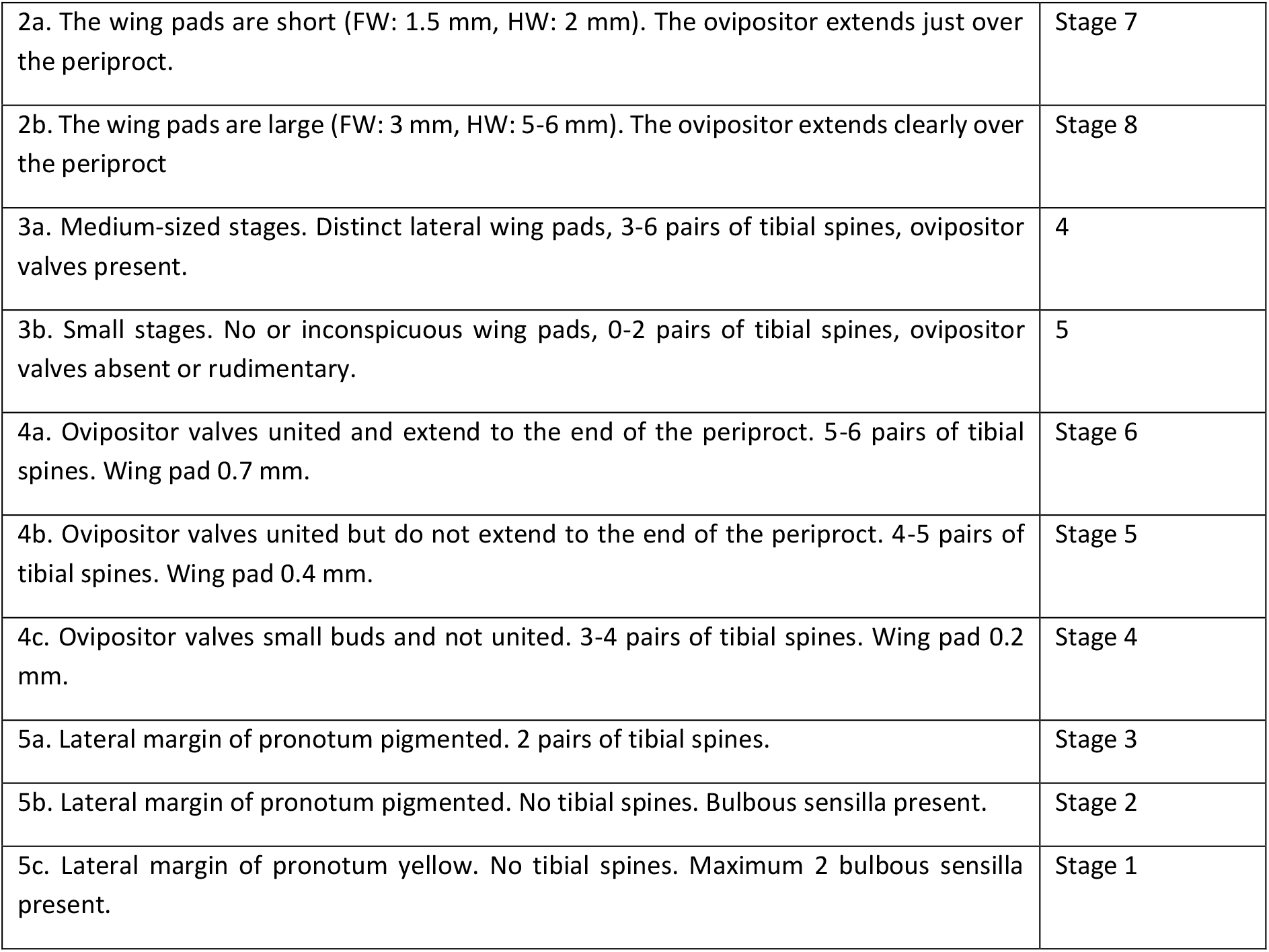
Identification key to the postembryonic stages.

Interestingly, the results show that the number of moults is reduced in favorable environments, which has been noted previously (Minelli and Fusco, 2013). However, both the proximate and ultimate cause for this phenomenon has been discussed to a lesser extent (Esperk et al., 2007). The process of moulting is energetically costly to the insect, so it seems contradictory that the number of moulting events is increased when energetic resources are scarce. It has been proposed that mortality of cricket nymphs is highest when the animal undergoes profound morpho-functional transformations (Zhemchuzhnikov and Knyazev, 2012). Since the turnover of wing buds coincides with the transition from the sixth to the seventh instar, additional moults in the sixth stage may indicate that the cricket prepares more carefully for this morphological transition in unfavorable conditions. From an evolutionary perspective, the cricket may use the variability of the sixth stage to fine-tune the duration of its life cycle so that it reaches adulthood in favorable conditions, i.e. when the ambient temperature is high enough for effective reproduction. Mechanistically, the increased number of moults may be linked to physiological constraints, for example the slower diffusion of oxygen in lower temperatures may force the insect to moult more often without having any adaptational relevance (Kivelä et al., 2016; Kivelä et al., 2018).

The postembryonic development of *G. bimaculatus* reflects the need for a conceptual change in the terminology commonly used to refer to postembryonic development in hemimetabolous insects (Minelli and Fusco, 2013). In terms of developmental biology, the term instar is misleading, as it is dependent on environmental conditions and does not correspond to morphological progression. Here the use of stages instead of instars is emphasised, so that postembryonic development encompasses 8 stages with 8-12 instars.

## 5. Conclusions

The postembryonic development of *G. bimaculatus* accentuates the need to reconsider the conceptual framework of arthropods, for a morphology-based stage does not always equate to one moult-based instar (Minelli and Fusco, 2013). With diverse molecular and genetic techniques at hand, *G. bimaculatus* offers an apt model system for understanding the mechanical underpinnings of this variability.

## Acknowledgements

This work was supported by Suomen Biologian Seura Vanamo ry, Suomen Hyönteistieteellinen Seura, and Entomologiska Föreningen i Helsingfors - Helsingin Hyönteistieteellinen Yhdistys (Pro gradu -apuraha).

